# Levels of sdRNAs in cytoplasm and their association with ribosomes are dependent upon stress conditions but independent from snoRNA expression

**DOI:** 10.1101/584367

**Authors:** Anna M. Mleczko, Piotr Machtel, M. Walkowiak, A. Wasilewska, K. Bąkowska-Żywicka

**Affiliations:** Institute of Bioorganic Chemistry Polish Academy of Sciences, Noskowskiego 12/14, 61-704 Poznań, Poland

**Keywords:** processed snoRNAs, snoRNA-derived small RNAs, sdRNAs, *Saccharomyces cerevisiae*, ribosome-associated noncoding RNA

## Abstract

In recent years, a number of small RNA molecules derived from snoRNAs have been observed. Findings concerning the functions of sdRNAs in cells are limited primarily to their involvement in microRNA pathways. However, similar molecules have been observed in *Saccharomyces cerevisiae*, which is an organism lacking miRNA machinery. Considering our previous observations, we examined the subcellular localization of sdRNAs in yeast. Our findings reveal that both sdRNAs and their precursors, snoRNAs, are present in the cytoplasm at levels dependent upon stress conditions. Moreover, both sdRNAs and snoRNAs interact with translating ribosomes in a stress-dependent manner. As a consequence of binding to ribosomes, yeast sdRNAs exhibit inhibitory activity on translation. However, observed levels of sdRNAs and snoRNAs in the cytoplasm and their association with ribosomes suggest independent regulation of these molecules by yet unknown factors.

## INTRODUCTION

Small nucleolar RNAs (snoRNAs) are noncoding RNAs that contribute to ribosome biogenesis and RNA splicing by modifying ribosomal RNA and spliceosomal RNAs, respectively. However, recently emerging evidence suggests that some snoRNAs have non-canonical functions in RNA editing, alternative splicing or maintenance of chromatin structure ^1–3^. The mechanistic details of these non-canonical functions are largely unknown. Moreover, it has recently become evident that mature, functional snoRNAs undergo processing to form stable short fragments, termed psnoRNAs, for processed snoRNAs ^4^ or sdRNAs for snoRNA-derived RNAs ^5^. The presence of processed forms of snoRNAs has been demonstrated in several organisms, including the primitive protozoan *Giardia lamblia* ^6^, Epstein-Barr virus ^7^, the budding yeast *Saccharomyces cerevisiae* ^8^, and mammals ^4^,^9–13^.

Concerning possible functions of small RNAs derived from snoRNAs (sdRNAs), their potential to regulate alternative splicing events ^4^ as well as their microRNA–like abilities, have been described in several organisms ^10^,^14^,^15^. Recently, it has been postulated that FUS-dependent sdRNAs in human cell lines might regulate gene expression, affecting transcript stability and translation ^16^. As the majority of miRNA-targeted, and thus translationally repressed, mRNAs are distributed in the cytoplasm, miRNA-like sdRNAs are expected to co-localize within the cytoplasm. Indeed, a subset of small RNAs derived from snoRNAs have been detected in the cytoplasm in *G. lamblia* ^14^ and humans ^10^; however, their nucleolar localization has been reported as well ^11^. Our recent work presented another possibility of sdRNA localization within the cytoplasm by association with ribosomes ^8^. These studies were performed in *S. cerevisiae*, an organism lacking miRNA machinery; hence, one might suspect a distinct role for non-miRNA sdRNAs in yeast.

It is now commonly accepted that full-length snoRNAs are not exclusively localized within the nucleus but are also present in the cytoplasm. Moreover, their cytoplasmic abundance is dynamically regulated in various stress conditions, such as oxidative stress ^17^, lipotoxic conditions ^18^ or heat shock ^19^. Knowledge concerning snoRNA expression regulation, however, is sparse. In 2002, it was shown for the first time that snoRNAs are involved in cancer development ^20^. Since that time, a growing body of evidence has emerged linking both full-length snoRNAs and their derivatives to oncogenesis (reviewed in ^21^). Considering both localization and the potential functions of full-length snoRNAs and sdRNAs, one might suspect that snoRNAs and their derivatives orchestrate responses to environmental stress outside of the nucleus. Surprisingly, studies aimed at revealing the abundance and subcellular localization of sdRNAs in stress have yet to be reported.

Yeast sdRNAs are observed in limited numbers during the sequencing of ribosome-associated small RNAs, with a maximum of 15 copies ^8^. However, their regulatory potential cannot be excluded, as recent studies demonstrated that even relatively small levels of ribosome-associated noncoding RNA (ranc_18mer, ∼27,000 molecules/cell) are sufficient to substantially influence global ribosome activity (∼200,000 ribosomes/cell) and regulate translation ^22^. We recently demonstrated that due to low abundance, conventional techniques, such as northern hybridization, are not sensitive enough to detect the full repertoire of cellular sdRNAs ^23^. Poor sensitivity and low throughput of hybridization-based technologies can be overcome by sensitive, amplification-based detection methods, such as stem–loop reverse transcription PCR (SL-RT-PCR), originally described by Chen et al. ^24^ and successfully implemented by our group for detection of yeast sdRNAs from as little as 50 ng of low molecular weight cellular RNA ^23^.

Therefore, to investigate the subcellular localization of both full-length snoRNAs and snoRNA-derived small RNAs in *S. cerevisiae*, we used this amplification-based method. To enable absolute quantification of RNAs, we implemented digital droplet PCR (ddPCR) technology. Our comprehensive analysis of sdRNA and snoRNA abundance and localization was performed under 12 different yeast growth conditions. Herein, we present the first evidence that snoRNA levels and the localization of sdRNAs within the cell, including association with ribosomes, are dependent upon stress conditions. As a consequence of sdRNA binding to ribosomes, an inhibition of *in vitro* and *in vivo* translation occurs. Moreover, for the first time, we present experimental data suggesting that both the expression and ribosome association of two types of related RNAs, namely, snoRNA-derived small RNAs and full-length snoRNAs, are independent from each other during multiple growth conditions.

## MATERIALS AND METHODS

### snoRNA-derived small RNAs

Three snoRNAs (snR67, snR83 and snR128) and 3 corresponding sdRNAs (sdR67, sdR83 and sdR128) were chosen for analysis based on the highest read coverage observed in ribosome-associated small RNA sequencing in *S. cerevisiae* ^8^. Sequences of sdRNAs are shown in Table 1. Localization of sdRNAs within predicted snoRNA secondary structure is shown in Figure 1.

**Table 1.**
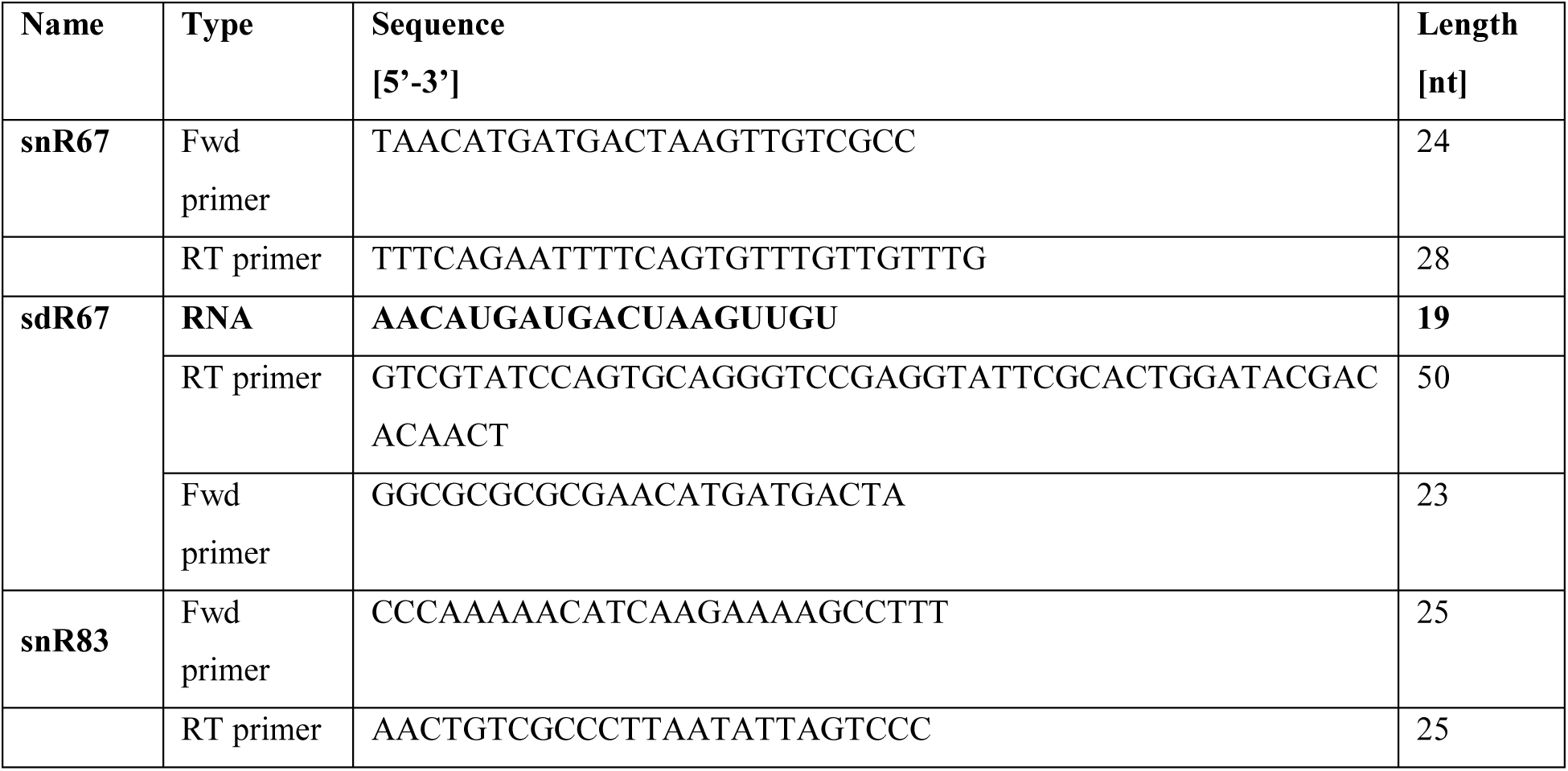
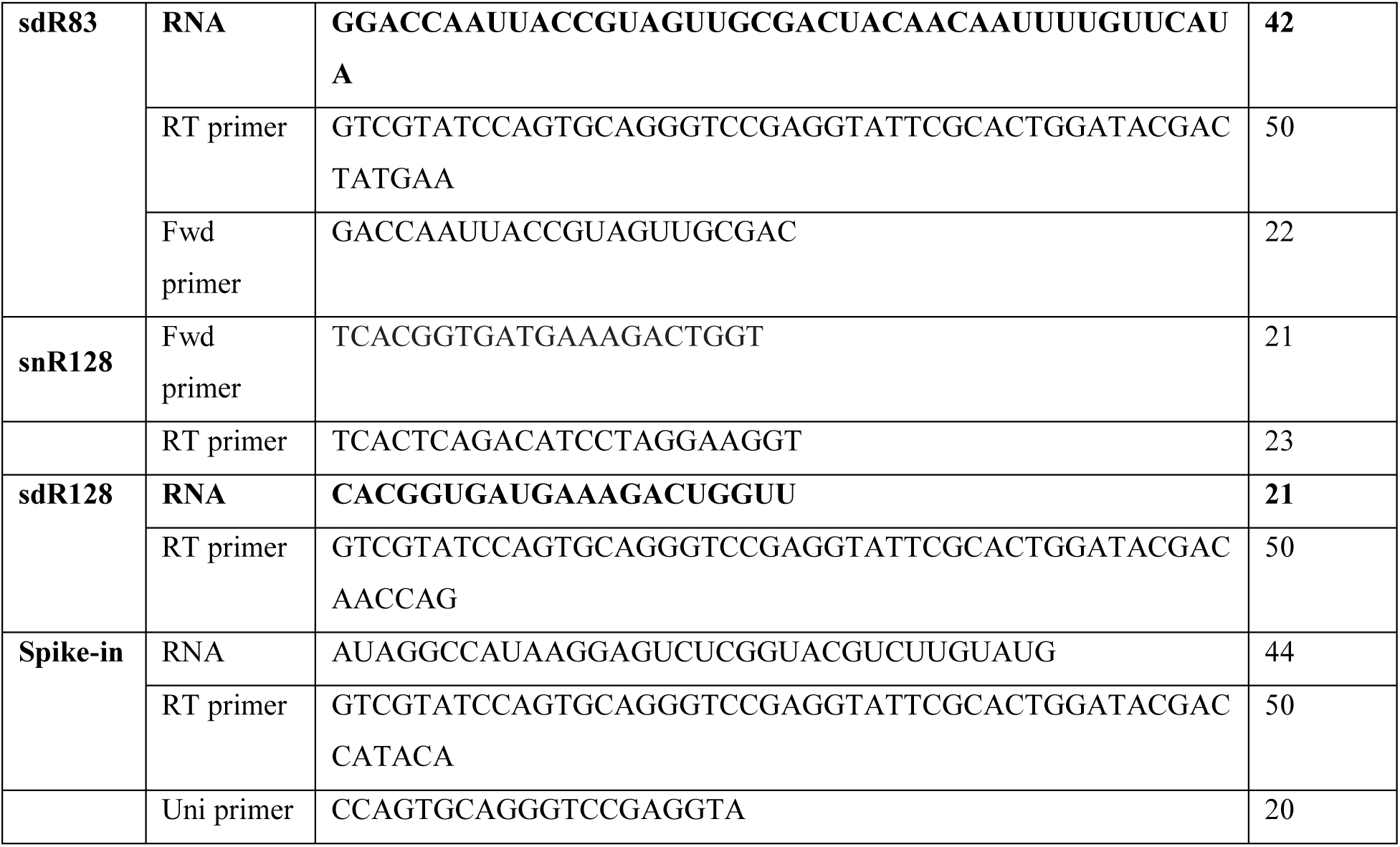
Oligonucleotide sequences used in this study. sdRNA sequences are presented in bold. RT primer – primer used for reverse transcription; Fwd primer – forward ddPCR primer; Uni primer – universal reverse ddPCR primer.

**Figure 1.**
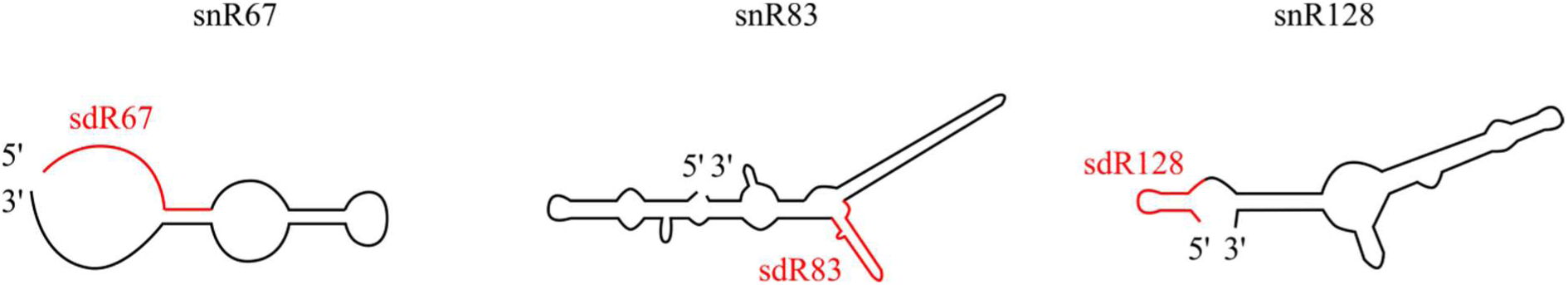
Localization of sdRNAs within predicted snoRNAs secondary structures.

### Strain and growth conditions

*Saccharomyces cerevisiae* strain BY4741 was grown in YPD medium with 2% glucose at 30°C. Environmental stress was induced as previously described ^25^ in two biological replicates. Briefly, cells were grown to mid-log phase, and stress conditions were applied for 15 min. Next, cells were pelleted and stored at −20°C. Stress conditions were as follows: heat shock (37°C), cold shock (15°C), high salt conditions (1 M NaCl), high pH conditions (pH 7.9), low pH conditions (pH 4.0), UV exposure (120 J/m^2^ UV), hyperosmotic shock (1 M sorbitol), hypoosmotic shock (cells grown to mid-log phase in YPD supplemented with 1 M sorbitol were transferred to YPD without sorbitol), amino acid starvation, sugar starvation, and anaerobic and normal growth.

### Yeast lysates and ribosome preparation

Yeast ribosomes were prepared as previously described ^26^. Briefly, cell pellets were washed with ice-cold water and resuspended in buffer (10 mM MgCl2, 100 mM KCl, 50 mM Tris/HCl, pH 7.5, 0.4 mM PMSF) at 4°C. Equal volumes of glass beads (400 μm in diameter) were added, and cells were broken using 8 pulses of vortexing (30 sec each) punctuated by cooling on ice. Cell debris was precipitated at 11,300 × g for 2 min at 4°C. Lysate was further clarified by centrifugation at 11,300 × g for 10 min at 4°C. After clarification, 1/10 of the total lysate volume was used to isolate total cellular RNA (S30). Subsequently, ribosomes were pelleted (P100) from lysates by centrifugation at 160,000 × g for 90 min at 4°C. Pelleted ribosomes (P100) and yeast lysate (S30) were mixed with TRI Reagent (MRC), flash frozen in liquid nitrogen and subjected to RNA isolation following the manufacturer’s instructions. Purity of P100 fractions and the quality of total cellular RNAs were verified using an Agilent Bioanalyzer 2100 with an RNA Nano 6000 kit.

### Reverse transcription

Stem-loop RT primers for sdRNA amplification (Table 1) were designed as previously described ^23^. Standard RT primers for snoRNA amplification were designed using the Primer3Plus tool. All reverse transcription reactions were performed in a multiplex manner. Reverse transcription reactions contained 10 or 100 ng RNA from P100 or S30 fractions, 50 nM of each stem-loop RT primer for sdRNAs and spike-in RNA, 50 nM of each standard RT primer for snoRNAs, 1×RT buffer, 0.25 mM of each dNTPs, 50 U SuperScript SSIII reverse transcriptase (Invitrogen), 5 U RiboLock RNase Inhibitor (Thermo Scientific), 10 mM DTT and 500 fM spike-in RNA (Table 1) as a normalizer. Twenty-microlitre reactions were incubated in a Bio-Rad T100TM Thermocycler for 30 min at 16°C, followed by pulsed RT of 60 cycles at 30°C for 30 sec, 42°C for 30 sec, and 50°C for 1 sec.

### Digital droplet PCR (ddPCR)

Copy numbers of sdRNAs and snoRNAs were determined using the QX100™ Droplet Digital™ PCR system (Bio-Rad, Pleasanton, CA). The reaction mixture was composed of 10 μl of 2x QX200™ ddPCR™ EvaGreen Supermix, 200 nM specific forward and universal reverse primers (Table 1), and 1 μl cDNA.

### Electrophoretic mobility shift assay (EMSA)

EMSA experiments were performed as previously described ^27^. RNAs corresponding to sdRNA sequences were chemically synthetized by Future Synthesis. A control RNA oligomer (5’-CUUGAGAUGAUUGCUAUGAUAC-3’) with no sequence similarity to any of the *S. cerevisiae* snoRNA sequences was used for all experiments. Ten picomoles of 5′-[32P]-end-labelled synthetic sdRNA was incubated with 25 pmol yeast ribosomes as previously described 32. Band shifts were resolved on non-denaturing 8% polyacrylamide gels and visualized on phosphor – storage intensity screen (Fujifilm) overnight. Screens were scanned using Fujifilm Fluorescent Image Analyzer FLA – 5100.

### Translation of poly(U) templates *in vitro*

Translation of poly(U) templates was performed as described ^28^ using 5 A260 units of ribosomes, 25 mg poly(U), 100 mg soluble protein factors, 25 mg deacylated yeast tRNA and 0.3 nmol [3H]-phenylalanine. The reaction was performed at 30°C for 30 min. Products were precipitated in TCA, recovered on Whatman glass fibre GF/C filters and subjected to scintillation counting. *In vitro* translation assays were performed in triplicate. Reported values are corrected for control samples lacking ribosomes, which were typically 0.5% to 1% of the total probe counts applied. P-values were calculated using t-tests.

### Metabolic labeling

Yeast spheroplasts were prepared from a 50-ml culture grown to an OD600 of 0.8 by adding 350 U of zymolyase (Zymo Research) and incubating at 30°C for 25-30 min as previously described ^22^. Spheroplasts were combined with synthetic sdR128 and electroporated. For controls, translation was inhibited by adding 7.5 μg/μl cycloheximide to the spheroplasts. Electroporated spheroplasts were incubated at 30°C with 1 μl 35S-methionine (1 000 Ci/mmol, 10 mCi/ml) for 1 h. Labelled proteins were precipitated in TCA, recovered on Whatman glass fibre GF/C filters and subjected to scintillation counting. Metabolic labelling measurements were performed in triplicate, and standard error (SE) was calculated. Statistical significance was determined using t-test.

## RESULTS

### SL-RT-ddPCR method enables for detection of small RNA input amounts

To perform highly sensitive detection of low abundance sdRNAs in yeast, we employed pulsed reverse transcription using stem-loop primers followed by digital droplet PCR (SL-RT-ddPCR). The ddPCR system measures fluorescence intensities of droplets after completion of all thermal cycling. The copy number of target genes is determined based on the number of fluorescent-positive and -negative droplets in a sample well. ddPCR provides an absolute number of RNA copies present in the sample. To define the minimum number of sdRNA copies that can be detected using the pulsed SL-RT-ddPCR method, we spiked 0.5 pg of an exogenous synthetic RNA (37 nt in length, no sequence similarity to *S. cerevisiae* snoRNAs) to total RNA isolated from *S. cerevisiae*. Total RNA was subjected to the SL-RT method. Various dilutions of cDNA were amplified using ddPCR technology. These analyses demonstrated the ability of ddPCR to detect small RNA input levels, as low as 0.005 pg (Suppl. Fig. 1).

### snoRNA and sdRNA levels are dependent upon stress conditions but are independent from each other

Using ddPCR, we investigated the accumulation of individual snoRNAs and sdRNAs across different *S. cerevisiae* growth conditions. Spiked-in synthetic RNA was used as a reference for experiments. Absolute concentrations of spike-in reference RNA in different cDNA samples were uniformly distributed, with a mean value of 19,064 (±114) copies/μl. Therefore, we concluded that possible differences in snoRNA or sdRNA concentrations under particular stress conditions would be derived from their abundance and not from biases in experimental design.

All full-length snoRNAs were least abundant under low pH stress, and absolute concentrations were as follows: 10,360 copies/μl for snR67, 717 copies/μl for snR83 and 13,200 copies/μl for snR128 (Fig. 2A and Suppl. Fig. 2A). Except for this stress, where the observed snoRNA concentrations were markedly lower, absolute concentrations under the remaining stress conditions were in a range of 169,000 – 578,200 copies/μl for snR67, 182,400 – 463,200 copies/μl for snR83 and 333,900 – 1,072,000 copies/μl for snR128. Under optimal yeast growth conditions, snR128 was significantly more abundant (698,000 copies/μl) than snR67 and snR83 (422,700 copies/μl and 440,700 copies/μl, respectively).

**Figure 2.**
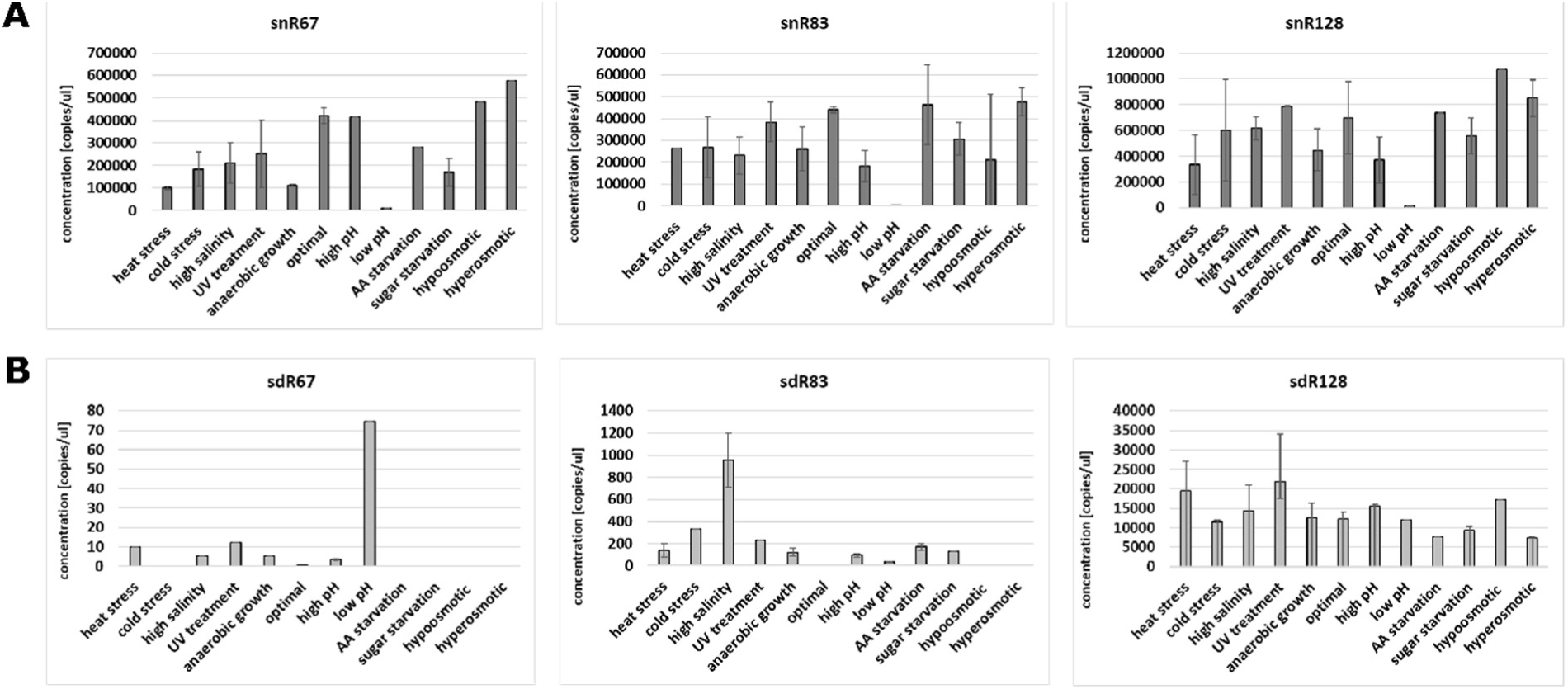
Quantitation of snoRNAs and sdRNAs within the total cellular RNA pool. Concentration (copies/microlitre) of snoRNAs (A) and sdRNAs (B). The mean and SE of two experiments are shown.

For sdRNAs, we clearly observed that sdR67 was present in the smallest levels compared to other sdRNAs, just above ddPCR detection level (Fig. 2B and Suppl. Fig. 2B). Its maximum concentration was noted under low pH conditions, at 74 copies/μl. sdR83 was moderately abundant compared to other two sdRNAs. The highest concentration of sdR83 was observed under high salinity conditions, and it reached 953 copies/μl. The most abundant of tested sdRNAs, sdR128, was equally distributed, with the most prominent concentrations under heat stress (19,470 copies/μl), UV shock (21,900 copies/μl) and hypoosmotic stress (17,220 copies/μl).

Since we observed clear differences in both snoRNA and sdRNA levels across different stress conditions, we next performed analysis of possible correlations between accumulation of these two molecules under particular types of stress (Fig. 3). We observed antagonistic changes under three stress conditions, namely, in low pH stress for snR67, high salinity for snR83 and heat stress for snR128. Under these three conditions, snoRNAs were significantly less abundant and sdRNAs were significantly more abundant. Apart from these observations, in most of the stress conditions, snoRNA levels did not correlate with sdRNA abundance. This observation suggests that differential accumulation of sdRNAs is not directly dependent upon the levels of individual snoRNAs under particular yeast growth conditions. This suggests possible stress-dependent regulation of sdRNA excision.

**Figure 3.**
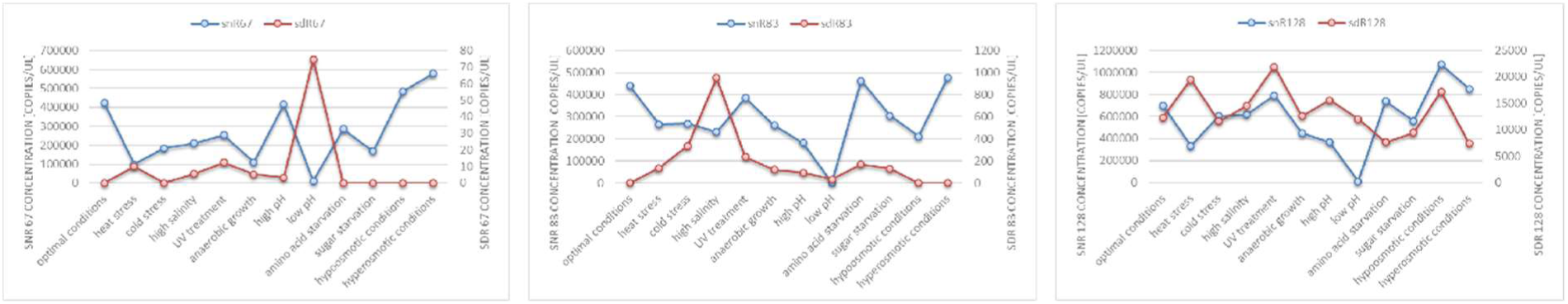
Differential accumulation of snoRNAs and sdRNAs in the cytoplasm. Values are means of replicates that are fully presented in Figure 2.

### snoRNAs and sdRNAs associate with ribosomes *in vivo* in a stress-dependent manner

Centrifugation of yeast lysates at 160,000 x g allowed us to isolate ribosomes and ribosome-associated RNAs (rancRNAs) from yeast cultivated under different growth conditions. For accurate quantification of ranc-snoRNAs and ranc-sdRNAs, we employed ddPCR technology. Spiked-in synthetic RNA was used as a control for experiments, as previously described for total cellular RNA pools. Absolute concentrations of spike-in reference RNA in different cDNA samples derived from ribosome-associated RNA pools were uniformly distributed, with a mean value of 7,573 (±11) copies/μl. Therefore, we concluded that possible differences in snoRNA or sdRNA concentrations in different rancRNA pools derive from their differential association with ribosomes and not from biases due to experimental design.

The first observation was that both full-length snoRNAs and sdRNAs strongly associate with mature cytoplasmic ribosomes (Fig. 4 and Suppl. Fig. 3). Moreover, analysis of snoRNA/ribosome and sdRNA/ribosome interactions in 12 different yeast growth conditions illustrate that this association is strongly stress-dependent. snoRNAs interact robustly with yeast ribosomes (Fig. 4A), exceeding association of sdRNAs over 800 times on average (Fig. 4B). In case of snR67, its highest concentration in ribosomal fractions was observed when ribosomes were isolated from yeast subjected to high pH conditions (292,400 copies/μl). The lowest snR67 concentration was noted in optimal conditions, as well as cold stress, and it oscillated approximately 2,800 copies/μl. snR83 was characterized by the lowest concentration among all three examined snoRNAs, with the maximum of 157,200 copies/μl in high pH conditions. Similarly, to snR67, cold stress caused the lowest accumulation of snR83 in the ribosomal fraction (22,100 copies/μl). The strongest association with ribosomes was observed for snR128, ranging from 57,300 copies/μl (hypoosmotic growth conditions) to 279,400 copies/μl (heat shock). In general, heat stress and high pH stress induced significant increases in ribosome-associated snoRNAs.

**Figure 4.**
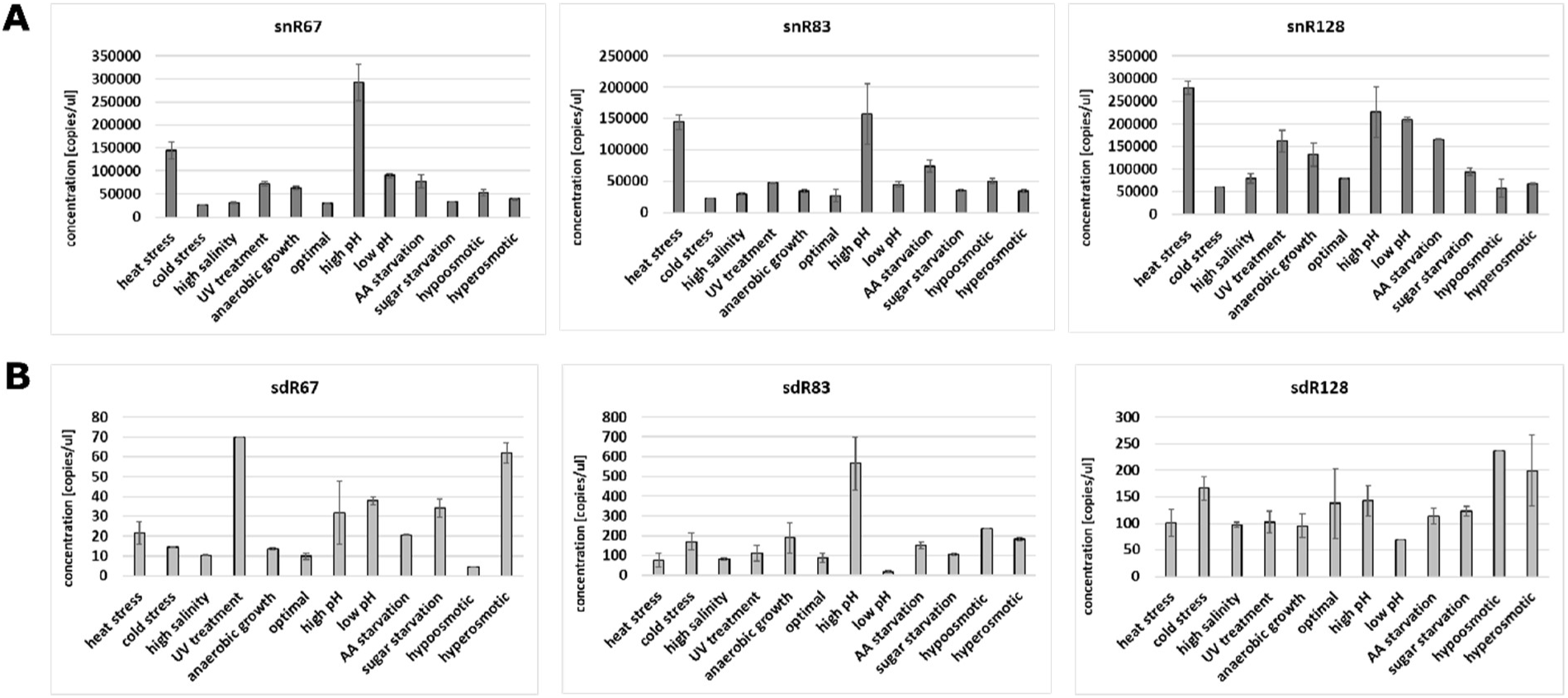
Quantitation of snoRNAs and sdRNAs within ribosome-associated RNAs. Concentration (copies/microlitre) of snoRNAs (A) and sdRNAs (B). The mean and SE of two experiments are shown.

Among investigated sdRNAs (Fig. 4B), sdR67 was the least abundant, reaching a maximum of 70 and 62 copies/μl after UV treatment and hyperosmotic stress conditions, respectively. The lowest accumulation of sdR67 in the ribosomal fraction was detected during hypoosmotic stress with 4.5 copies/μl. Generally, the amount of sdR83 was significantly higher in most stress conditions compared to sdR67. The highest concentration was observed during high pH conditions (565.5 copies/μl). In contrast, in low pH conditions, only 18.5 copies/μl of sdR83 were detected. For sdR128, a variety of growth conditions did not strongly affect its presence in the ribosome-associated RNA pool. Highest sdR28 accumulation was observed in hypo- and hyperosmotic conditions, reaching 236 and 199 copies/μl, respectively.

To elaborate on *in vivo* interactions with ribosomes, we compared the differential accumulation of both snoRNAs and sdRNAs in ribosome-associated RNA fractions (Fig. 5). In cases of high pH stress, both snR83 and its derivative, sdR83, were highly abundant. Except for this case, in the remaining stress conditions, snoRNA and sdRNA levels within ribosome-associated RNA pools were not well correlated. Such observation suggests that stress-dependent association of full-length snoRNAs and small sdRNAs with yeast ribosomes is independent.

**Figure 5.**
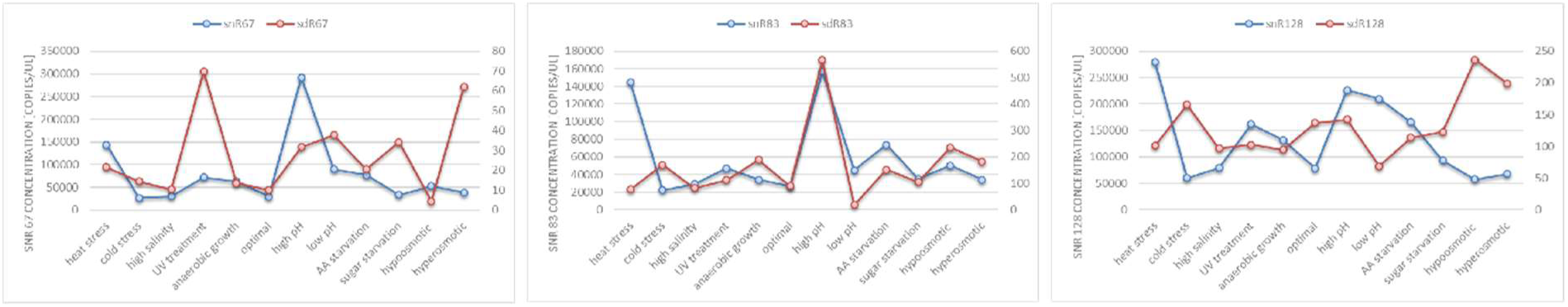
Differential accumulation of snoRNAs and sdRNAs in ribosomes. Values are means of replicates that are fully presented in Figure 4.

### sdRNAs interact with ribosomes *in vitro* in a stress-dependent manner and inhibit translation

To verify the observations that sdRNAs interact with yeast ribosomes in a stress-dependent manner, we performed electrophoretic mobility shift assay (EMSA) experiments. We grew yeast under 12 different conditions, isolated ribosomes and incubated them with synthetic radioactively labelled sdRNAs. Resulting mixtures were subjected to electrophoresis in polyacrylamide gels under native conditions. Formation of complexes between sdRNAs and ribosomal components occurred because migration of complexes was slower than corresponding free sdRNAs (Fig. 6).

**Figure 6.**
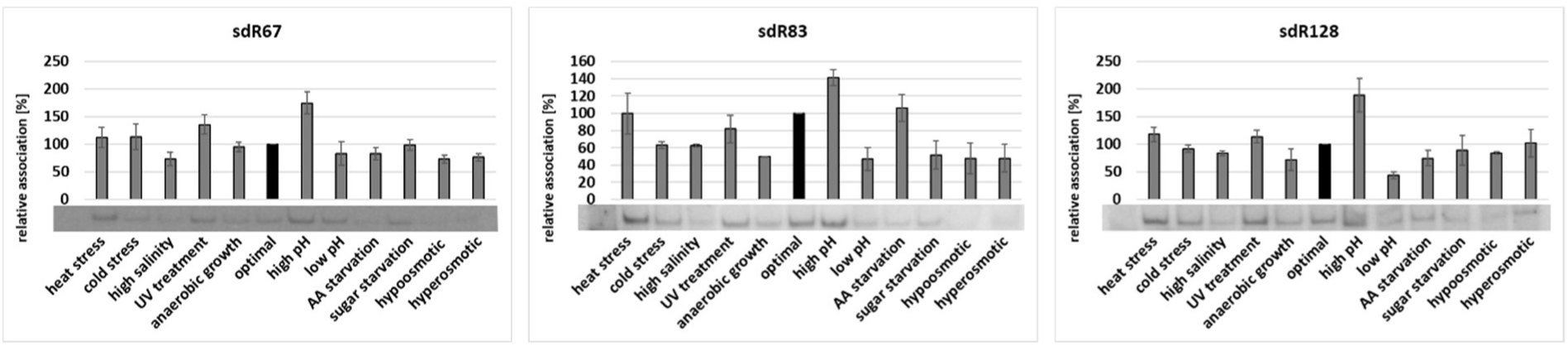
sdRNAs interact with *S. cerevisiae* ribosomes *in vitro*. sdRNA/ribosome associations in relation to optimal growth conditions (100%) is shown. The mean and SE of three experiments are shown. Representative sdRNA/ribosome complexes revealed by EMSA experiments are shown at the bottom of the x-axis.

All three investigated sdRNAs were most prone to interact with ribosomes isolated from yeast cultured under high pH stress. sdR67 and sdR128 binding to ribosomes was ∼2-fold more efficient than to ribosomes derived from yeast grown under optimal conditions. Except for high pH, under the overwhelming majority of stress conditions, sdRNAs were less prone to interact with ribosomes than under optimal growth conditions. Under high salinity, anaerobic growth, low pH or hypoosmotic stress, all sdRNAs formed reduced complexes compared to controls. In heat stress, we observed almost no difference in a number of complexes formed between stress and optimal conditions.

The observation that sdRNAs bind to ribosomes in a stress-dependent manner led to speculation of their potential function as regulatory ncRNAs during protein biosynthesis. To clarify this, we set up an *in vitro* translation system for *S. cerevisiae* grown under optimal conditions and quantified the number of synthesized polypeptides in the presence or absence of sdRNA. For this experiment, we chose sdR128, which associated with yeast ribosomes under multiple stress conditions did not differ significantly and was prominent under optimal yeast growth conditions (Fig. 4B). We observed a clear dose-dependent reduction in translation efficiency of poly(Phe) mRNA (Fig. 7A). We did not observe any increase in the inhibitory potential of sdR128 when dose was increased to over 100 pmol per translation reaction. To study the role of ribosome-associated sdRNAs in a physiological context, we performed metabolic-labelling experiments using yeast spheroplasts. We measured ^35^S-Met incorporation into newly synthetized proteins in the presence and absence of sdR128. We observed that yeast sdR128 decreased translational efficiency *in vivo* (Fig. 7B). Inhibition efficiency was in a range similar to the well-known ribosome-targeting antibiotic cycloheximide.

**Figure 7.**
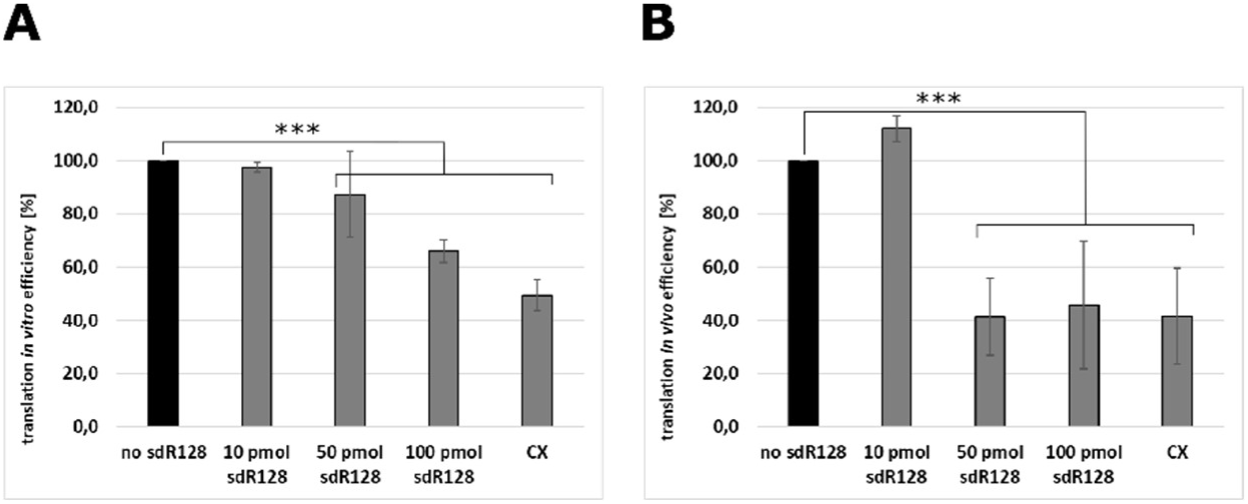
*S. cerevisiae* snoRNA-derived RNAs inhibit protein biosynthesis. Cycloheximide (CX) served as a control translational inhibitor. The mean and SE of three experiments are shown. (A) Dose-dependent effects of sdR128 on yeast poly(Phe) synthesis in an isolated in vitro translation system. Translational efficiency is shown as a percentage [%] of activity of the control experiment without sdR128. *** p-value <0.005 (B) Incorporation of 35S-methionine into the translatome of yeast spheroplasts is presented as translation in vivo efficiency [%]. The efficiency of metabolic labelling in the absence of sdR128 was set at 100%, and spheroplasts harbouring synthetic sdR128 were compared to this value. *** p-value <0.005

## DISCUSSION

In a recent study, we revealed that snoRNAs and sdRNAs in *Saccharomyces cerevisiae* are present in the cytoplasm, where they associate with ribosomes. Herein, we show that binding of *S. cerevisiae* sdRNA to ribosomes influences protein biosynthesis. Moreover, both cytoplasmic abundance and ribosome association of snoRNAs and sdRNAs are strongly dependent upon stress conditions. For the first time, we have shown that snoRNA and sdRNA levels in the cytoplasm and their association with ribosomes are independent from each other.

Quantitative determination of low levels of small RNAs remains challenging. Because low abundant sdRNAs are not detectable using standard methods, such as northern blot hybridization ^23^, we decided to employ an optimized stem-loop reverse transcription (SL-RT) followed by ddPCR. We have shown that SL-RT-ddPCR method is sensitive enough to detect as low as several copies/μl of sdRNAs in *S. cerevisiae*. However, in order to enable reverse transcription, stem-loop primers have to be complementary to the last six nucleotides at the 3’ end of individual sdRNAs. As a consequence it is not possible to distinguish small sdRNAs derived from 3’ ends from their longer precursor snoRNAs during the analysis of ddPCR results. Thus, we were able to detect only sdRNAs derived from the 5’ end (sdR128, sdR67) or from the middle part of snoRNAs (sdR83). For precise detection of RNAs derived from 3’ end of their precursors, one additional size selection step in the procedure would be needed. One possibility is to separate small and long RNAs on gels and elute them from the precise spaces on the gel. A prominent disadvantage of such size – selection is the fact that after elution we have two pools of isolated RNAs, which might differ in quality and quantity.

It has been already shown that snoRNAs play crucial roles in adaptation to stress conditions ^21^. In yeast, U3 snoRNA (snR17) is upregulated during heat shock, amino acid starvation and sugar starvation and downregulated under high salinity and hyperosmotic conditions ^29^. Interestingly, the expression pattern of snoRNAs studied herein do not resemble those of snR17. In Chinese hamster ovary (CHO) cells U14 snoRNA, which corresponds to yeast snR128, is strongly upregulated during heat shock ^19^. In yeast, however, we observed almost 2-fold downregulation of snR128 during heat stress. Such observations suggest that different snoRNAs possess distinct expression patterns during stress responses, which might be related to their functions.

Despite the fact that dysregulation of snoRNAs is involved in adaptation to stress conditions, only a few publications have reported the roles of sdRNAs in tumour development, which may be considered a stress for human cells ^21^,^30^,^31^. Expression analysis of sdRNAs has been performed in several types of cancer with the conclusion that accumulation of sdRNAs is associated with malignant transformation and that increased global production or accumulation of sdRNAs is already occurring in the early stages of cancer. Surprisingly, there is no data reporting sdRNA levels in canonical stress conditions. To our knowledge, differential expression of sdRNAs has not previously been reported. Here, we report for the first time that sdRNAs in *S. cerevisiae* are present under a wide repertoire of growth conditions, though in some cases in limited amounts. So far, sdRNAs have been reported to localize in the cytoplasm in organisms where they act within microRNA pathways ^6^,^10^,^12–14^. Here, we present new data demonstrating cytoplasmatic localization of sdRNAs in an organism that lacks miRNA pathways.

The presence of both sdRNAs and snoRNAs in ribosome-associated RNAs indicates the possible existence of a novel, yet to be discovered stress-dependent translation regulation mechanism. The data presented herein strongly suggest that specific interactions between yeast ribosomes and sdR128 downregulate translational activity during optimal growth conditions. In this aspect, sdR128 could be classified as an example of ribosome-associated noncoding RNAs (rancRNAs), next to an mRNA exon-derived 18-residue-long ncRNA ^22^ and tRNA-derived fragments ^26^ previously described by our lab.

We demonstrated interactions between sdRNAs and yeast ribosomes both *in vivo* and *in vitro*. Some degree of similarity between *in vitro* binding of synthetized sdRNAs to ribosomes in EMSA experiments and in cellular levels of ribosome-associated sdRNAs measured with ddPCR have been observed (Suppl. Fig. 4). We believe an imperfect correlation between these two methods might be a consequence of much lower sensitivity of EMSA compared to ddPCR or/and the requirement for additional factors for ribosome association in living cells.

Moreover, for the first time, we present data showing that snoRNAs and sdRNAs are differentially abundant in both the cytoplasm and in ribosomes. Such observations indicate that sdRNAs and snoRNAs might perform distinct cellular functions in response to stress conditions, probably during translation regulation, as shown here for sdR128 under optimal conditions. These observations strongly support the hypothesis for separate roles of both snoRNAs and sdRNAs during stress conditions.

## Acknowledgements

This work was supported by the National Science Centre, Poland [UMO-2014/13/D/NZ1/00061 to K.B.Ż.] and [UMO-2017/27/B/NZ1/01416 to K.B.Ż]. The work was also supported by the Polish Ministry of Science and Higher Education, under the KNOW programme.

## Authors contribution

K.B.Ż. planned experiments. A.M.M, P.M., A.W and M.W performed experiments. K.B.Ż., A.M.M. and P.M. analysed the data and wrote the paper.

## Supplementary materials

**Supplementary Figure 1.**
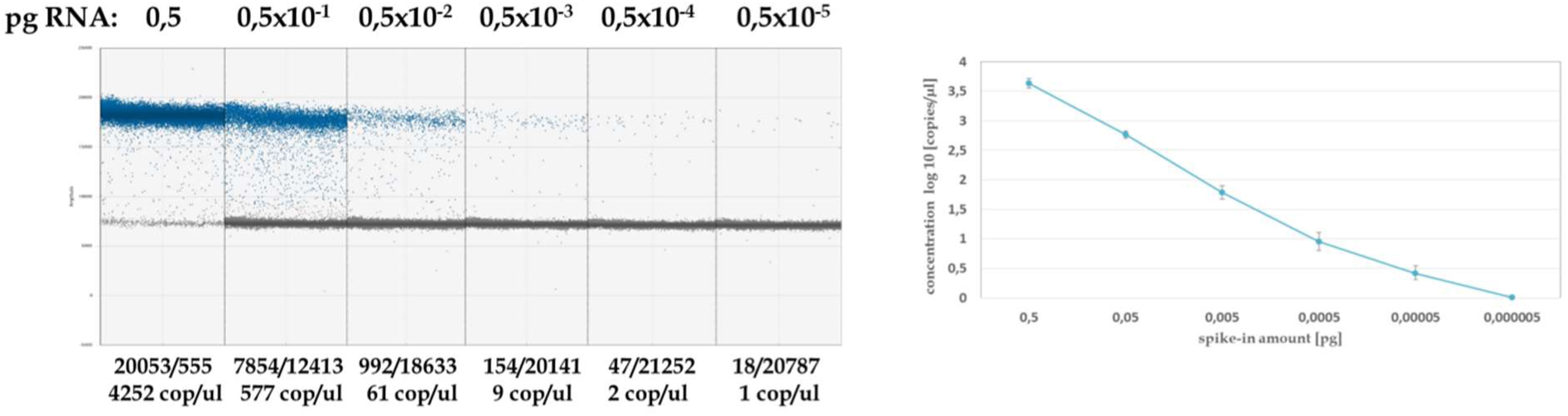
Detection and quantitation of synthetic spike-in RNA. A) Fluorescence amplitude of different concentrations of spike-in RNA. Levels of input RNA are reported at the top, whereas the number of positive/negative droplets and the concentration (copies/microlitre) are reported on the bottom of each panel. B) The relationship between calculated copies/microlitre and input RNA.

**Supplementary Figure 2.**
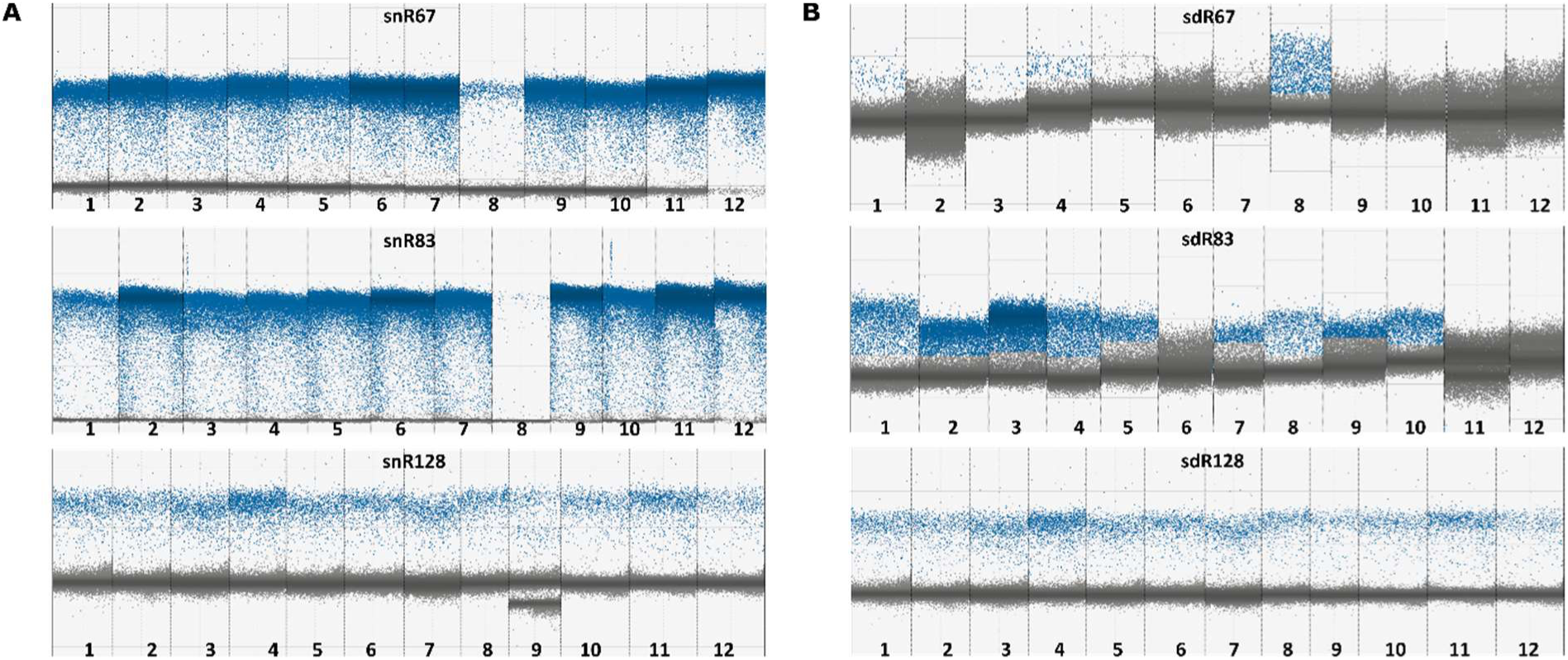
Detection of snoRNAs and sdRNAs within total cellular RNA pools. Representative fluorescence amplitude of snoRNAs (A) and sdRNAs (B). Positive droplets are in shown in blue, negative droplets in grey. Slot numbers correspond to stress conditions as follows: 1 – heat stress, 2 – cold stress, 3 – high salinity, 4 – UV treatment, 5 – anaerobic growth, 6 – optimal conditions, 7 – high pH, 8 – low pH, 9 – AA starvation, 10 – sugar starvation, 11 – hypoosmotic conditions, and 12 – hyperosmotic conditions.

**Supplementary Figure 3.**
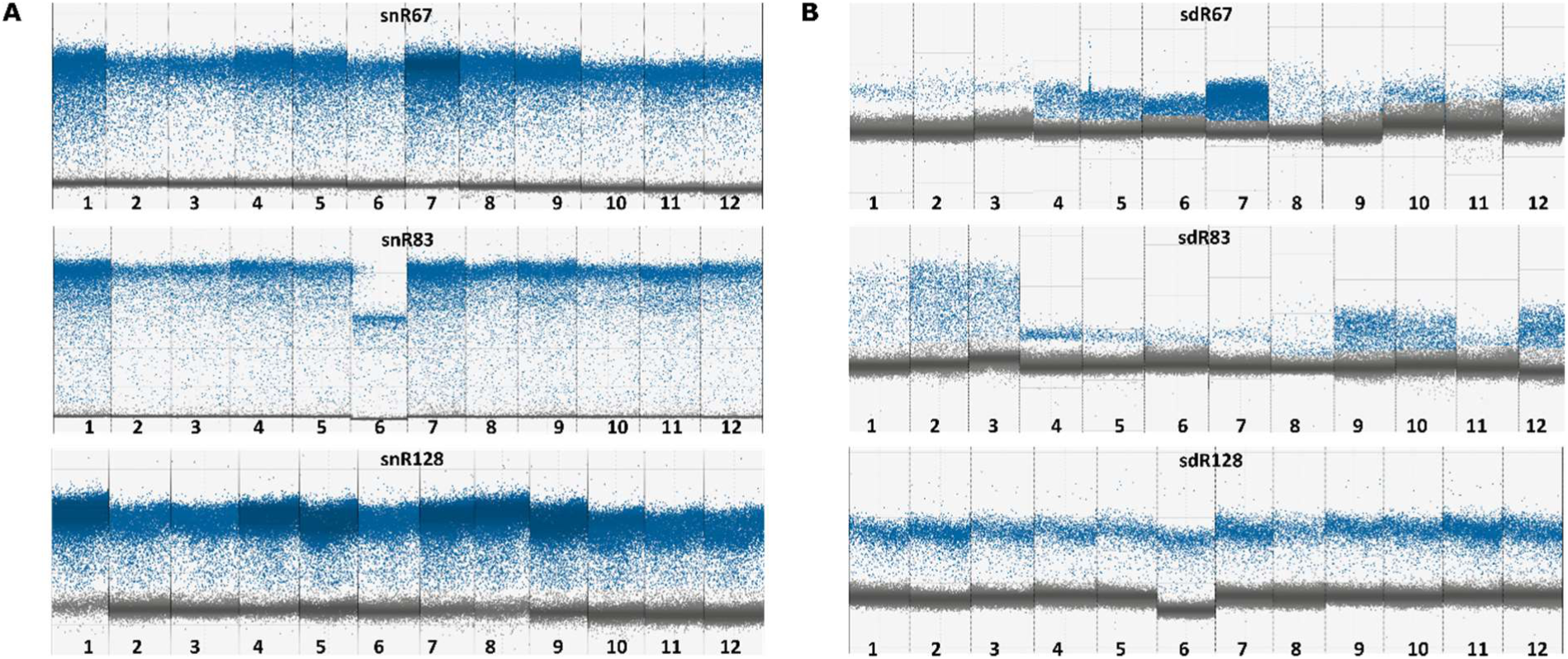
Detection of snoRNAs and sdRNAs within ribosome-associated RNAs. Representative fluorescence amplitude of snoRNAs (A) and sdRNAs (B). Positive droplets are in shown blue, negative droplets in grey. Slot numbers correspond to stress conditions as follows: 1 – heat stress, 2 – cold stress, 3 – high salinity, 4 – UV treatment, 5 – anaerobic growth, 6 – optimal conditions, 7 – high pH, 8 – low pH, 9 – AA starvation, 10 – sugar starvation, 11 – hypoosmotic conditions, and 12 – hyperosmotic conditions.

**Supplementary Figure 4.**
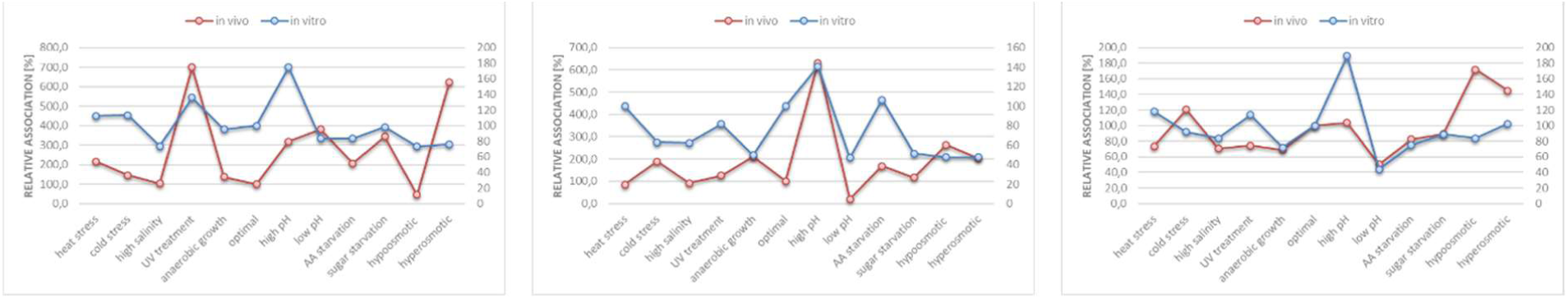
Differential association of sdRNAs with ribosomes *in vivo* and *in vitro*. Relative association [%] was calculated in relation to values under optimal conditions (100%).

